# Comparison of the cortical hierarchy between macaque monkeys and mice based on cell-type specific microcircuits

**DOI:** 10.1101/2024.10.31.621258

**Authors:** Mélissa Glatigny, Françoise Geffroy, Timo van Kerkoerle

## Abstract

The primate neocortex contains a hierarchy of cortical areas, with feedforward connections running from lower to higher levels, and feedback connections running in the opposite direction. The relative hierarchical position of cortical areas has been well established by retrograde tracing studies that allow to determine whether type of output from different source areas is predominantly feedforward or feedback. This method can not determine whether the cortico-cortical input within target areas is predominantly feedforward or feedback. We here make use of cell-type specific microcircuits to provide an intrinsic measure of the strength of feedforward versus feedback processing within cortical areas in macaque monkeys and mice. This allows a more complete map of the cortical hierarchy of different species, and therefore a more direct cross-species comparison. Parvalbumin-expressing interneurons were used as a marker of feedforward processing and calretinin-expressing interneurons as a marker of feedback processing. We found steep gradients in the distributions of these two interneuron types across macaque monkey cortical areas, indicating a deep cortical hierarchy where early visual areas are dominated by feedforward neural circuits and higher cortical areas become dominated by top-recurrent circuits. In contrast, the gradient in interneuron distribution across mouse cortical areas was limited, indicating a shallow cortical hierarchy and remaining dominated by feedforward neural circuits. This implies that the mouse cortex is comparable to early visual areas in primates, remaining dominated by bottom-up input. While primates are unique in having a deep cortical hierarchy that allows neural processing in higher cortical areas to become predominantly internal.

**Significance statement:** The cortex in primates is known to contain a hierarchy of cortical areas, where bottom-up processing allows sensory information to get more and more compressed towards higher cortical areas, and where top-down processing allows lower areas to remain informed about the internal goals of the animal. However, it has been challenging to directly compare cortical hierarchies between animal species. Here, we developed an intrinsic measure of the strength of bottom-up and top-down processing within cortical areas, based on the density of different types of interneurons within areas. Or results indicate that the mouse cortical hierarchy is shallow and is driven by bottom-up processing, while the primate cortical hierarchy is deep and slowly becomes dominated by internal processing towards the top.

## Introduction

A defining architectural principle of the primate neocortex is that it contains a functional hierarchy of distinct cortical areas, with large areas towards the bottom of the hierarchy that encode elementary sensory features, and smaller areas towards the top of the hierarchy that encode more complex and abstract information (1–4). This functional hierarchy has an anatomical substrate with a consistent laminar organization in the connectivity pattern between areas : neurons in superficial layers (2 and 3) send feedforward connections mainly targeting layer 4 of the next level, while neurons in the deep layers (5 and 6) send feedback connections mainly targeting layer 1 of the previous level of the hierarchy (**Figure 1**) (5–7).

**Figure 1.**
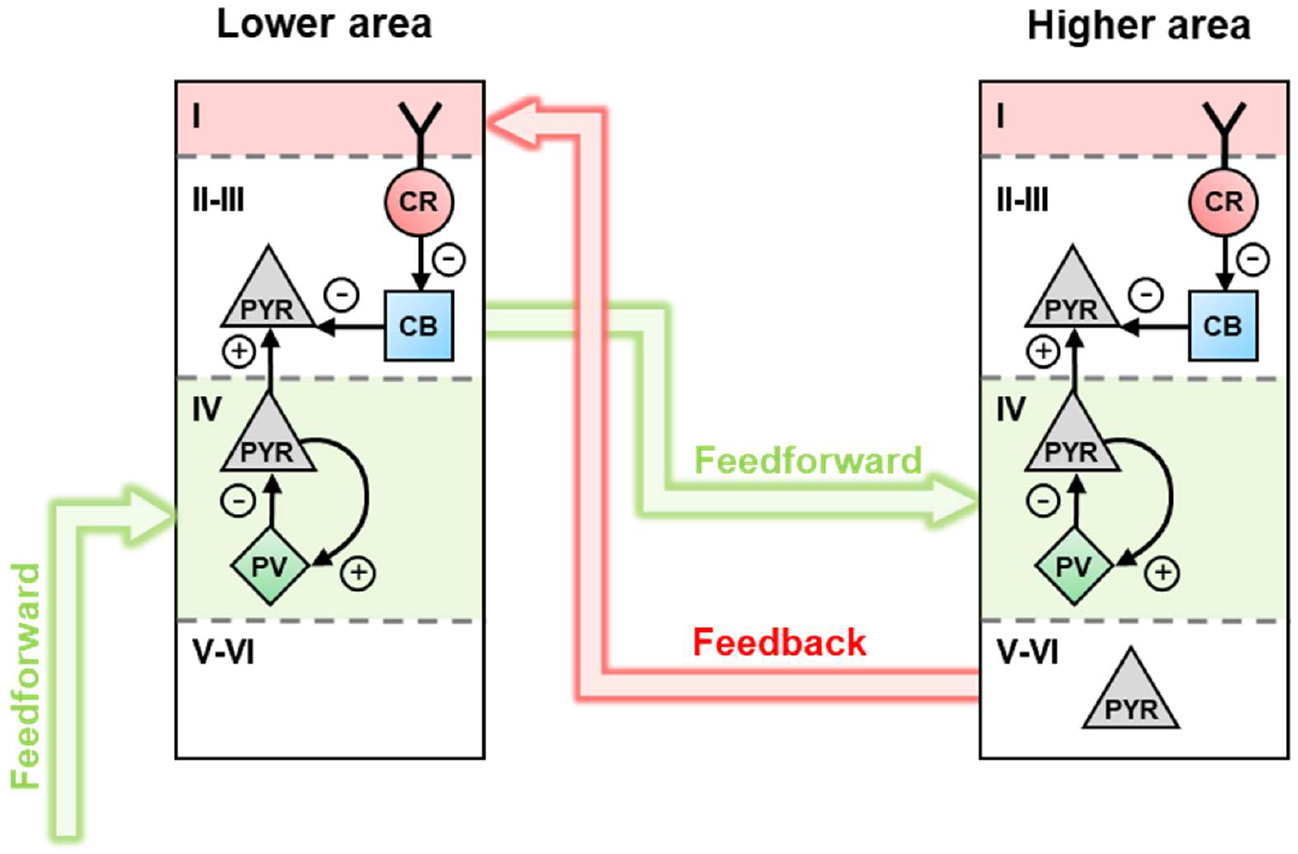
Schematic illustration of the interneuron-specific microcircuits involved in feedforward versus feedback processing. Feedforward connections (green) predominantly target the circuit between parvalbumin interneurons (PV) and pyramidal cells (PYR) in layer 4. In contrast, feedback connections (red) terminate predominantly in layer 1, targeting calretinin (CR) or VIP interneurons which disinhibit PYR cells through calbindin (CB) or somatostatin (SST) interneurons. See text for further explanation.

This laminar organization of the connectivity pattern can be used to quantify the hierarchical position of different cortical areas relative to each other. This is based on the injection of a retrograde tracer in a target area and mapping the laminar position of neural cell bodies within different source areas (7). Within a given source area, the more cell bodies there are in the superficial layers relative to the deep layers, the more this area is thought to be providing feedforward connections to the target area, and the other way around for feedback connections. This approach has been extensively applied to macaque monkeys and mice to establish the relative hierarchical position of cortical areas within these species (1, 7, 8). However, this method cannot determine how input from source areas is integrated within target areas, it therefore cannot provide an intrinsic measure of the position of different areas within the cortical hierarchy. This has led to at least two important debates about the organization of the cortex.

First, how is the relative strength of feedforward versus feedback processing distributed across the cortex? It is commonly argued that feedback connections are more numerous than feedforward connections (7, 9–12), and that V1 takes a special place as it is all the way at the bottom of the hierarchy, receiving a cascade of top-down input (13–15). This interpretation seems reasonable considering that V1 only receives ∼1% of its external input from the lateral geniculate nucleus (LGN), and ∼90% from higher cortical areas, relative to the total number of neurons in different source areas (7). In contrast, others have argued that internal amplification circuits within V1 allow it’s neural processing to remain dominated by the feedforward drive from the LGN, which is thought to be required for a reliable transmission of bottom-up input to the rest of the cortex (16–20).

Second, how should cortical hierarchies be compared between animal species, in particular when they strongly differ in brain size and number of cortical areas? For example, the cortex in mice has been suggested to be much less hierarchical compared to primates (8, 21, 22). This has been interpreted as implying that mouse cortex is dominated by recurrent interactions, with a strong integration of different sensory modalities (8). This would imply that neural processing within mouse cortex is similar to primate prefrontal cortex where a hierarchy is difficult to discern as well (4). In contrast, others have proposed that the mouse cortex is similar in architecture to early sensory areas in primates, and that an extensive association cortex is unique to primates (23–25).

These opposing views remain difficult to reconcile based on currently available data. An intrinsic measure of the strength of feedforward versus feedback input within each cortical area would provide a more complete view on the position of an area within the cortical area, allowing a more direct comparison of the cortical hierarchies across species. One way to do that would be by applying anterograde tracers in a specific source area and trying to map the laminar profile of axon boutons in different target areas (26). However, this is highly challenging in large brains and has there been done by only very few primate studies and restricted to very few cortical areas (5, 27). Moreover, to understand the impact that these boutons have on the target region would require trans-synaptic labelling techniques to map the type of neurons that are targeted by the axon boutons, which is technically even more challenging for animals with large brains.

Interestingly, it has recently been shown that two distinct interneuron-specific microcircuits are involved in feedforward versus feedback processing streams (**Figure 1**). While parvalbumin-expressing interneurons (PV) are selectively involved in fast feedforward inhibition (28–34), vasoactive intestinal peptide (VIP) expressing interneurons are implicated in top-down driven disinhibition via somatostatin (SST) expressing interneurons (33–43). This implies that PV and VIP interneurons could be used as anatomical markers of the strength of feedforward versus feedback input within a given cortical area. And measuring the relative density of PV and VIP interneurons across different cortical areas could provide a new approach to measure the intrinsic strength of feedforward versus feedback processing across the cortical hierarchy. This relatively straight forward method could thereby help to resolve the two debates described above.

The recent functional studies showing the involvement of PV and VIP in feedforward versus feedback processing are largely restricted to mice, based on genetic tools which are not easily accessible for use in primates. Recent anatomical work using transcriptomics has found genetic differences in cell-types between mice and monkeys (44–47). Still, the overall pattern of separate cell types being involved in feedforward versus feedback processing also holds true for macaque monkeys (28, 29). Classic post-mortem staining studies in primate cortex have shown that PV interneurons are selectively expressed in cortical layers that are targeted by feedforward connection (**Figure 1**) (28, 48, 49). And also in primates, PV interneurons selectively target the somas of pyramidal cells in these feedforward input layers (28), which would be appropriate for fast feedforward inhibition (29, 33, 50). Post mortem staining for VIP is challenging, but transcriptomics studies have shown that calretinin (CR) is nearly exclusively co-expresses with VIP interneurons in mice and macaque monkey cortex (44–47). And like VIP interneurons in mice, CR interneurons in primates are expressed in the cortical layers that are targeted by feedforward connection (**Figure 1**) (28). Crucially, also CR has been implicated in disinhibition by preferentially targeting other classes of inhibitory neurons in the superficial layers of both primate and rodent cortex (29, 51–53). And such a disinhibitory circuit would be appropriate to implement a relatively slow gain control (40), corresponding to the neural mechanisms that has been found in top-down processing (e.g. selective attention) in monkeys and humans (50, 54–56).

Some anatomical work has been already done to map the density of cortical interneurons in different animal species (57–60), but no dataset is available that uses the same method to map the relevant interneurons across both animal species and cortical regions. Here, we made a direct comparison between the density of PV and CR interneurons across the visual hierarchy in mice and macaque monkeys, including a range of areas within the ventrolateral prefrontal cortex (vlPFC) in macaque monkeys. We found a steep gradient of interneuron distribution in the hierarchy of macaque monkey cortex, suggesting that the neural microcircuits involved in feedforward processing are dominating early visual areas while feedback processing becomes dominant within vlPFC. In contrast, mouse cortex shows only a weak gradient in interneuron distribution and the complete cortex remains dominated by the neural circuits involved in feedforward processing.

## Results

We tested a large number of cell-type specific antibodies in mouse and macaque monkey cortex and found that staining worked reliably for PV, CR and NeuN (**Figure 2A**), while many others such as for VIP and SST worked well in mice, but not in macaque monkey cortex (**Table S1**), being either not sensitive or selective. This is in line with results from other labs (personal communication with Jonathan Ting from the Allen brain Institute). To allow a direct comparison between the strength of feedforward versus feedback processing across the visual hierarchy, we used triple immunolabeling against PV, CR as well as NeuN. This allows an expression of the two types of interneurons relative to the total number of neurons within each slice, which is relevant because of the substantial heterogeneity in the density across even neighboring slices. We restricted ourselves to areas of which the relative hierarchical position is well established based on anatomical and physiological studies (1, 4, 7, 21, 22, 61) (**Figure 2B**) (see the paragraph on “Brain areas” in the method section). In total, we collected an extensive number of neurons (**Table 1**) in 2 macaque monkeys and 3 mice (**Table S2**).

**Table 1.**
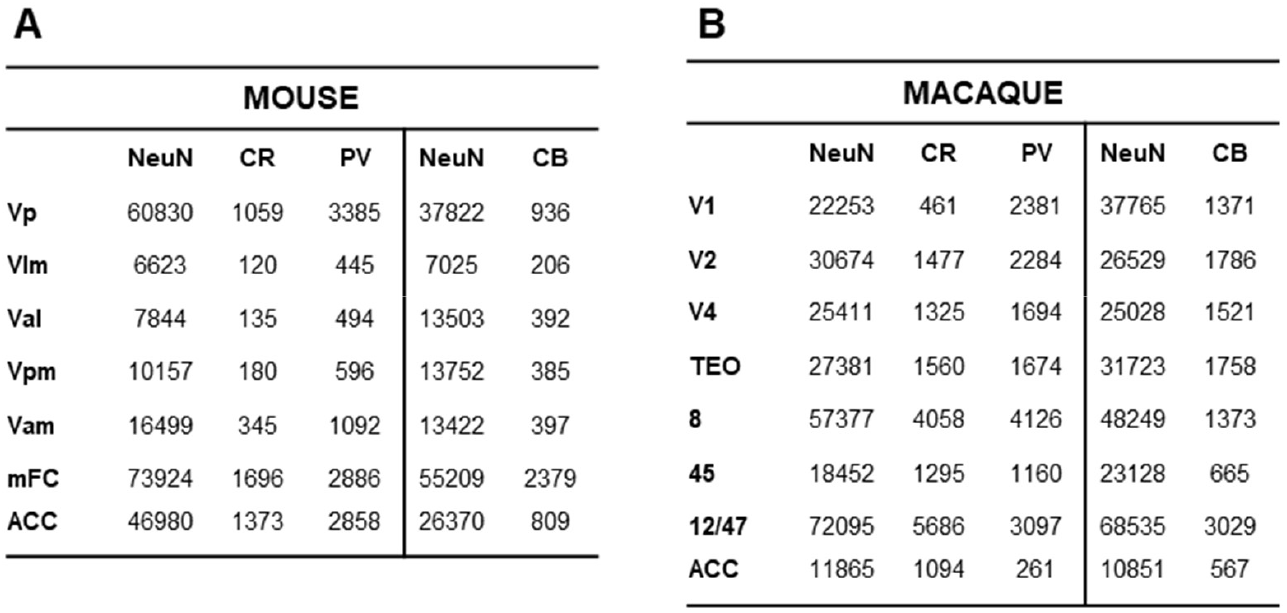
Total number of counted neurons in macaque monkey and mouse cortex. Number of total neurons and calcium-binding proteins counted for each cortical area in A) mice (n=3) and B) macaque monkeys (n=2).

| MOUSE |  |  |  |  |  |
| --- | --- | --- | --- | --- | --- |
|  | NeuN | CR | PV | NeuN | CB |
| Vp | 60830 | 1059 | 3385 | 37822 | 936 |
| VIm | 6623 | 120 | 445 | 7025 | 206 |
| Val | 7844 | 135 | 494 | 13503 | 392 |
| Vpm | 10157 | 180 | 596 | 13752 | 385 |
| Vam | 16499 | 345 | 1092 | 13422 | 397 |
| mFC | 73924 | 1696 | 2886 | 55209 | 2379 |
| ACC | 46980 | 1373 | 2858 | 26370 | 809 |

| MACAQUE |  |  |  |  |  |
| --- | --- | --- | --- | --- | --- |
|  | NeuN | CR | PV | NeuN | CB |
| V1 | 22253 | 461 | 2381 | 37765 | 1371 |
| V2 | 30674 | 1477 | 2284 | 26529 | 1786 |
| V4 | 25411 | 1325 | 1694 | 25028 | 1521 |
| TEO | 27381 | 1560 | 1674 | 31723 | 1758 |
| 8 | 57377 | 4058 | 4126 | 48249 | 1373 |
| 45 | 18452 | 1295 | 1160 | 23128 | 665 |
| 12/47 | 72095 | 5686 | 3097 | 68535 | 3029 |
| ACC | 11865 | 1094 | 261 | 10851 | 567 |

**Figure 2.**
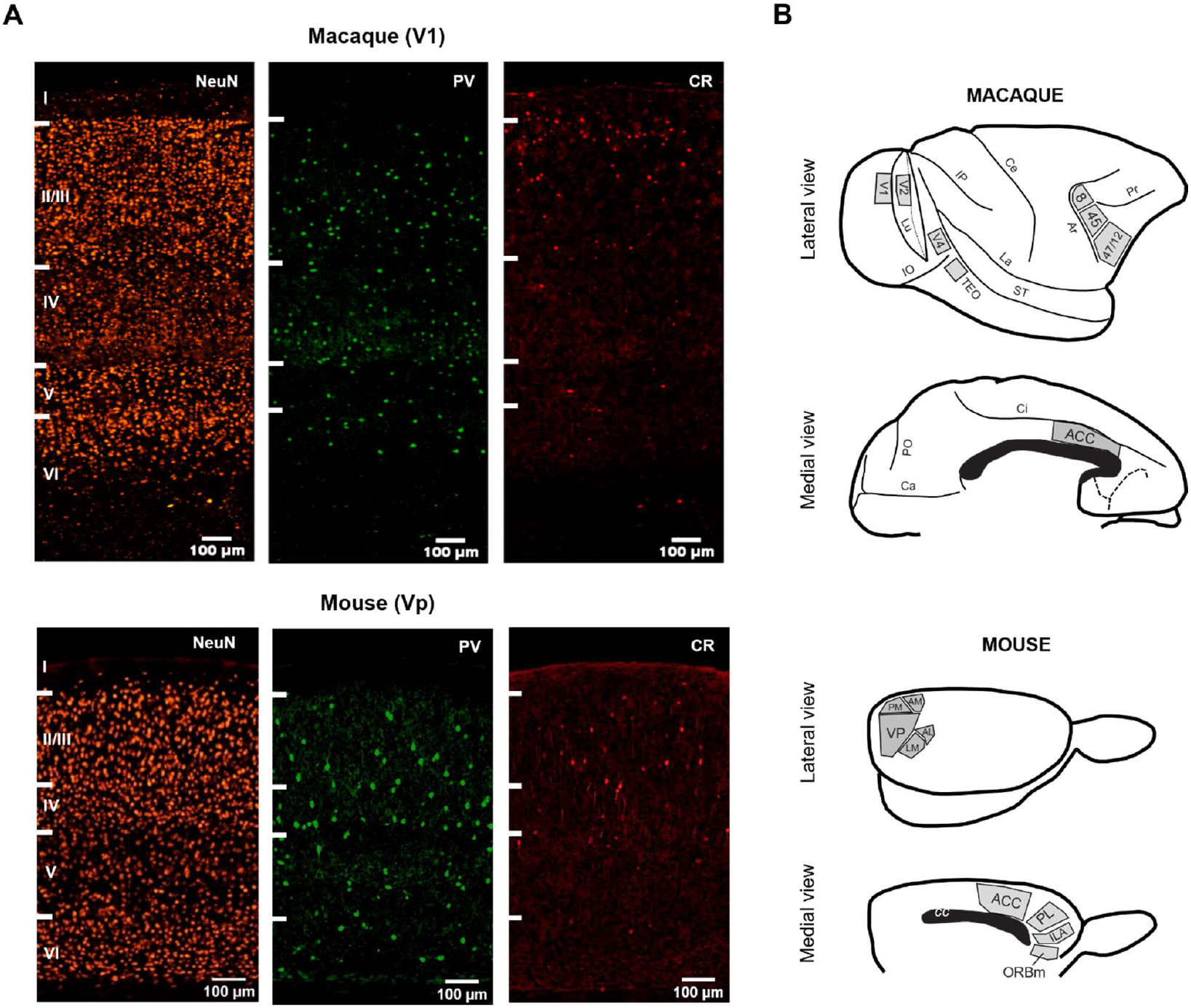
Experimental paradigm. (A) Representative examples of the laminar distribution for neurons (NeuN, orange), parvalbumin (PV, green) and calretinin (CR, red) immunoreactive cells in the primary visual cortex, (B) Schematic of the cortical areas that were investigated. Lu, lunate sulcus; IO, inferior occipital sulcus; ST, superior temporal sulcus; La, lateral fissure; IP, intraparietal sulcus; Ce, central sulcus; Ar, arcuate sulcus; Pr, principal sulcus; Ci, cingulate sulcus; Ca, calcarine fissure; PO, parieto-occipital sulcus.

Across the macaque monkey cortical hierarchy we found that the relative density of PV and CR interneurons showed steep gradients, where the density of PV decreased and the density of CR interneurons increased towards higher cortical areas (t-test of the slope of the regression line in visual cortex for PV interneurons, t(40)= 5.94, *P <* 0.0001; and for CR interneurons, t(40)= 4.20, *P <* 0.0005; t-test of the slope of the regression line in frontal cortex for PV interneurons, t(40)= 7.57, *P <* 0.0001; and for CR interneurons, t(50)= 3.06, *P <* 0.005) (**Figure 3A**). PV was the dominant interneuron in early visual areas (t-test, t(82)= 6.45, *P <* 0.0001), while CR became the dominant interneuron in the vlPFC (t-test, t(102)= 6.35, *P <* 0.0001) (**Figure S1**). This suggests that the neural circuits involved in feedforward processing are dominant towards the bottom of the hierarchy while recurrent processing dominates the top hierarchy. The normalized difference between CR and PV could thereby allow a measure of the position of each area within the cortical hierarchy, with −1 representing the bottom of the hierarchy and +1 representing the top of the hierarchy (**Figure 3B**). This indicates an orderly progression of the different cortical areas, which stretches from V1 being dominated by feedforward processing to ACC being dominated by recurrent processing.

**Figure 3.**
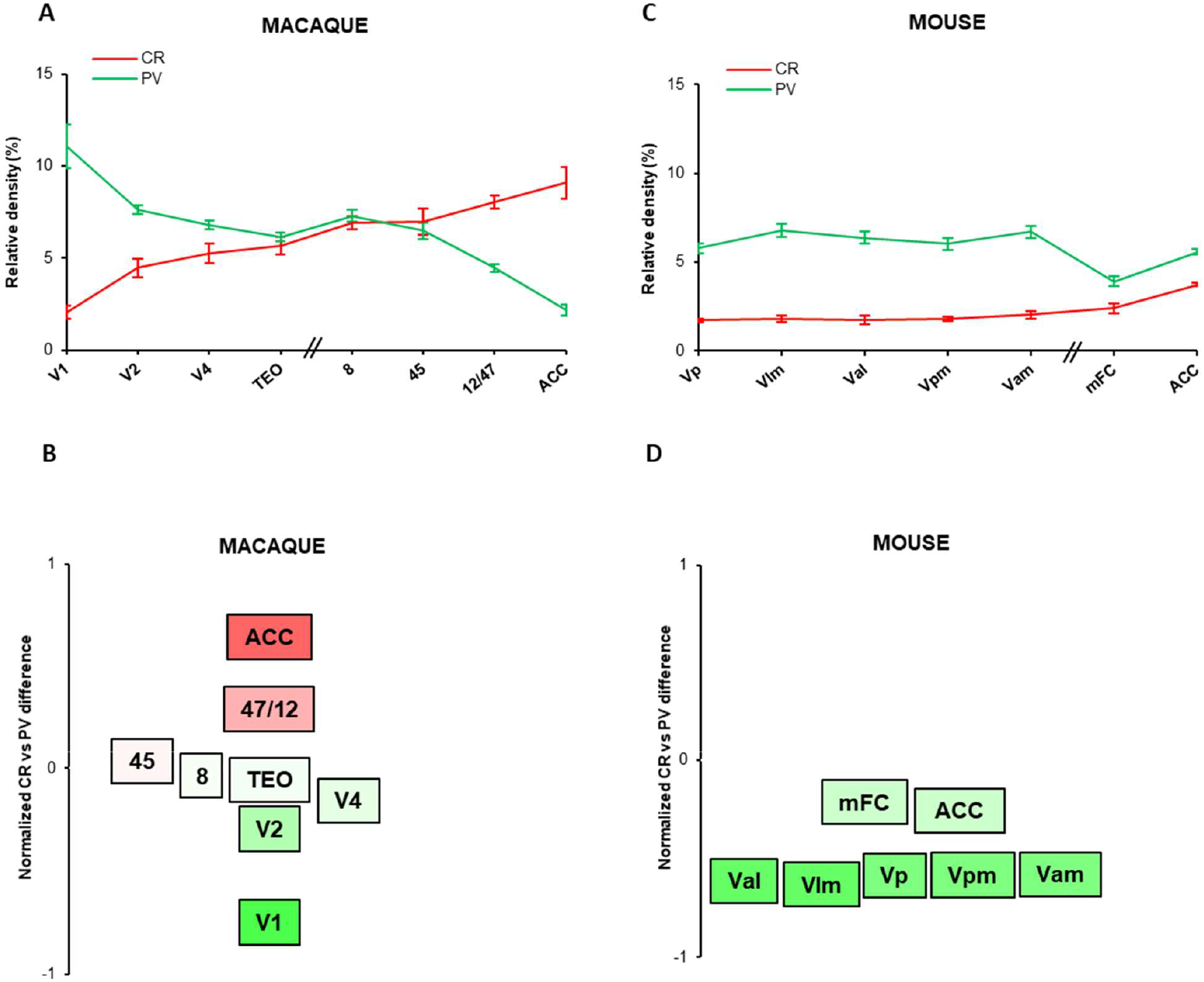
Interneuron distribution in the cortical hierarchy of macaque monkeys and mice. Density of two interneuron cell types, parvalbumin (PV, green) and calretinin (CR, red) relative to the total neuronal population (as estimated with NeuN staining) across different cortical regions for A) macaque monkeys and C) mice. Error bars indicate standard error of the mean. Normalized difference of CR and PV of the different cortical areas in B) macaque monkeys and D) mice, calculated as (CR-PV)/(PV+CR). The colors correspond to the value of the normalized difference, with bright red corresponding to 1, bright green corresponding to −1 and white corresponding to 0.

Strikingly, a gradient in the relative interneuron density was absent in mouse visual cortex (t-test of the slope of the regression line in visual cortex, p>0.5) (**Figure 3C**). Only a decrease of PV and increase of CR interneurons was found in mouse frontal relative to visual cortex (t-test for PV interneurons, t(323)= 8.27, *P <* 0.0001; t-test for CR interneurons, t(213)= 7.04, *P <* 0.0001), but PV remained more abundant than CR interneurons across cortical areas in mice (t-test for visual areas, t(139)= 26.5, *P <* 0.0001; t-test for frontal area, t(254)= 8.36, *P <* 0.0001) (**Figure S1**). This suggests that mouse cortex only contains a very shallow hierarchy, which remains dominated by feedforward processing, comparable to early visual areas in macaque monkeys (**Figure 3D**).

If PV and CR interneurons play a similar role in macaque monkeys and mice, their laminar density profiles should be expected to remain similar, as feedforward and feedback input target similar layers in macaque monkeys and in mice (**Figure 4**). In both macaque monkeys and mice, feedback connections mainly target layer 1 while feedforward connections mainly target layer 4 (5–7, 22, 38) (**Figure 1**). And the neurons that dominate a specific layer can be expected to be the dominant recipient of the respective afferent input (62). It is relevant to note that in mice, the output from the LGN is not restricted to the primary visual cortex but also targets higher visual areas (63, 64), in line with the interpretation that a cortical hierarchy is limited in mouse visual cortex. To investigate the laminar profile of PV and CR interneurons we calculated the density of these interneurons relative to the total number of neurons per layer, and averaged the densities across the different areas in the visual cortex and the frontal cortex in monkeys and mice. We separated the agranular frontal cortex which lacks a layer 4 (ACC in monkeys and ACC and mFC in mice), from the granular frontal cortex (i.e. areas 8, 45, 47/12 in the macaque vlPFC). The results for the granular frontal cortex are limited to monkeys as a mouse homologue is lacking for this region (25, 65). In all regions, both species showed a clear feedback laminar profile for CR density, peaking in layer 1 and dropping down to layer 2 and 3 (**Figure 4**). In contrast, PV density showed a feedforward laminar profile, peaking in the middle layers. In mouse visual cortex, PV density did not differ significantly between layer 4 and 5. The difference in the laminar profile between PV and CR is also visible when the densities are normalized relative to the number of neurons across a cortical column (instead of per layer), although differences in the density within layer 1 becomes very difficult to judge as the absolute number of neurons in this layer is very low (**Figure S2**).

**Figure 4.**
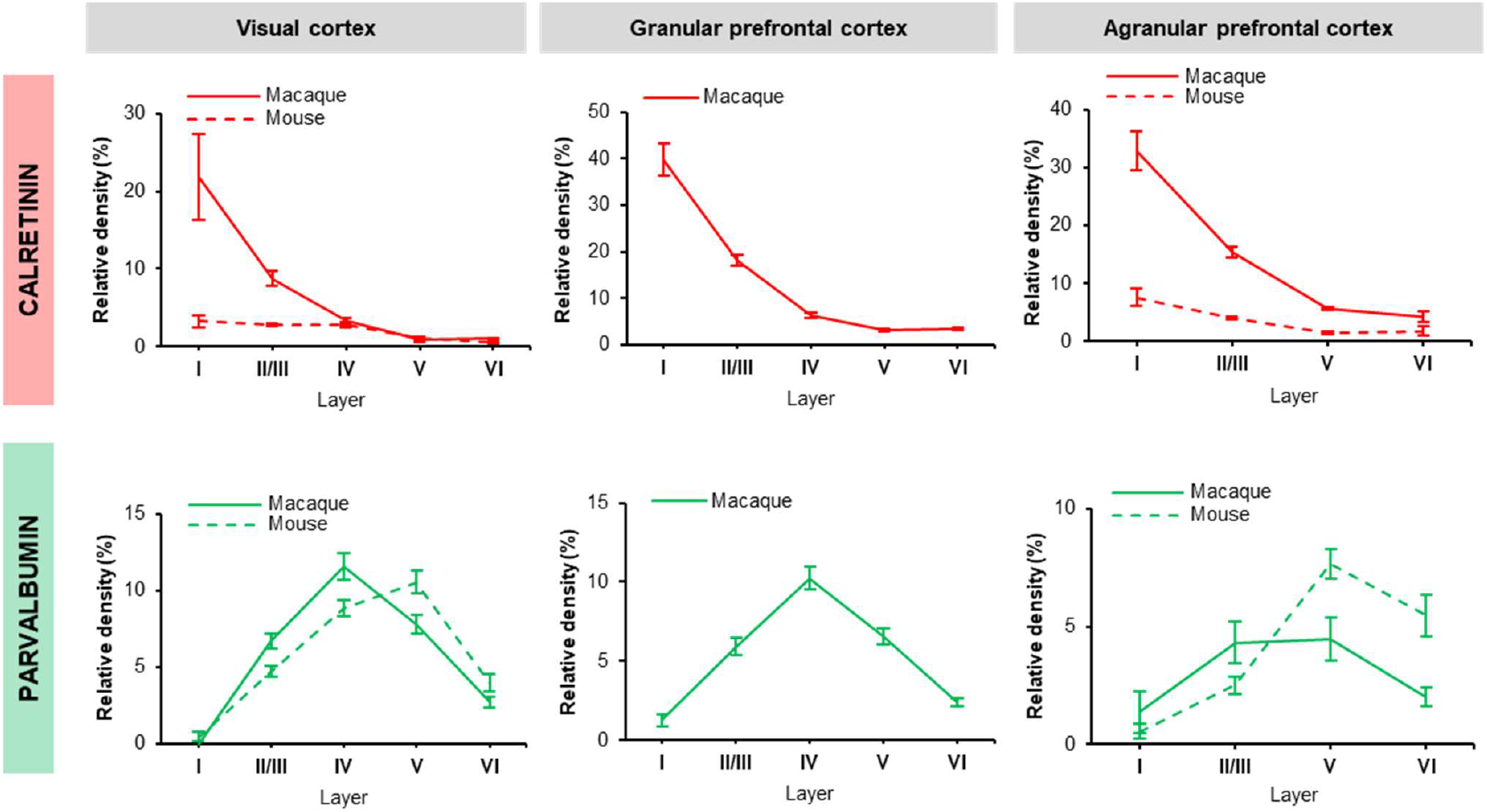
Laminar distribution of parvalbumin and calretinin interneurons. Densities of parvalbumin (bottom row) and calretinin (top row) interneurons relative to the total neuronal population within each laminar compartment of the visual (left column), granular frontal (middle column), and agranular frontal cortex (right column) in monkeys (continuous line) and mice (dashed line). Mice lack a granular frontal cortex, so for this region data is only shown for macaque monkeys. Error bars indicate standard error of the mean.

In separate brain sections, we also stained against CB interneurons, together with NeuN. CB interneurons show substantial genetic overlap with SST interneurons (44–47). CB could thereby provide an additional marker for feedback processing, as SST interneurons are indirectly involved in the feedback-driven disinhibitory circuit as well (**Figure 1**) (33–36, 38–40, 43). However, SST cells are also known to be involved in surround inhibition and therefore in horizontal processing (32–34), making a functional interpretation less straight forward. Furthermore, CB is expressed more widely in other cell types as well, even in pyramidal cells, with potential differences between species and cortical areas (45, 47, 66). Still, we stained against CB interneurons and NeuN in separate brain sections for the two macaque monkeys and in 3 separate mice. The distribution of CB followed the distribution of CR interneurons, in mice and monkeys, although with a drop in density between visual cortex and frontal cortex in monkeys (**Figure S3**). This confirms the gradient of recurrent circuits across the visual hierarchy as shown more directly with CR interneuron density. These results for the distribution of PV, CR and CB interneuron densities across cortical areas were consistent across individuals (**Figure S4**).

The sum of all calcium-binding protein labelled interneurons (PV, CR and CB) was higher in macaque monkeys compared to mice (t-test for visual areas, t(413)= 9.61, *P <* 0.0001; t-test for frontal areas, t(219)= 8.96, *P <* 0.0001), in line with previous results (67, 68), but was relatively stable across areas (grey line in **Figure S3**). Furthermore, the pattern in the density of the 3 types of interneurons when averaged across all visual areas was similar between monkeys and mice, with PV cells being most prevalent (t-test for PV versus CB interneurons for monkeys t(71)= 4.80, *P <* 0.0001; t-test for PV versus CB interneurons for mice, t(158)= 19.8, *P <* 0.0001) and CR neurons being least prevalent (t-test for CR versus CB interneurons for monkeys t(74)= 2.62, *P <* 0.01; t-test for CR versus CB interneurons for mice, t(182)= 8.13, *P <* 0.0001)(**Figure S1**), in line with previous results (60). While in frontal cortex, specifically the density of CR interneurons is strongly elevated in macaque monkeys compared to mice (t-test for CR interneurons in mice versus monkeys t(73)= 17.2, *P <* 0.0001; t-test for CB interneurons in mice versus monkeys t(86)= 0.19, *P*=0.42; t-test for PV interneurons in mice versus monkeys t(186)= 2.64, *P <* 0.01), matching previous studies as well (57–59, 69, 70).

## Discussion

We found marked differences in the distributions of interneuron densities within the cortical hierarchies of macaque monkeys and mice. A steep gradient is present in the primate cortical hierarchy, with the density of PV interneurons decreasing and CR increasing towards higher cortical areas, and with CR becoming prevalent within the vlPFC. As PV is thought to be involved in bottom-up processing and CR in top-down processing, this indicates that there is a steep gradient in the type of neural processing taking place throughout the primate cortical hierarchy, with bottom-up processing dominating early visual areas and top-down processing dominating higher cortical areas. This is fits well with the view that it is these higher order or associatory areas that receive multiple streams of bottom-up input which need to be dynamically selected based on behaviorally relevant goals that are kept in working memory (41, 71). We did not find a similar gradient in interneuron distribution in the mouse visual cortex, although we did find a similar decrease of PV and increase in CR interneurons from the visual to the frontal cortex, but with PV remaining more prevalent than CR interneurons. This suggests that in contrast to macaque monkeys cortex, mouse cortex contains a shallow hierarchy and remains dominated by bottom-up processing.

Although no previous study directly compared the relative distribution of PV and CR across the cortical hierarchy in macaque monkeys and rodents, a similar result can be observed in the normalized ratio of PV and CR when previous literature is combined (**Figure S5**) (48, 57, 58, 60, 66, 69, 70, 72, 73). A strong gradient can be consistently observed in primate cortex, with CR becoming dominant within higher cortical areas. In contrast, a gradient is absent or very weak in rodent cortex, with PV interneurons remaining prevalent across rodent cortex. Importantly, this indicates that these findings are consistent across differences in histological techniques, as well as consistent across a large number of individual animals.

Transcriptomic studies have found some genetic differences in cortical cell types between primates and rodents (44, 45), which could imply that the same cell type might serve different functions in different species. However, the distribution of RNA across the brain as measured with transcriptomics does not translate directly to protein levels as measured with antibody staining, and currently available transcriptomic data might therefore not be suited to investigate gradients in the distribution of different cell types across the cortex (47). Importantly, we did find that the laminar profiles of PV and CR are very similar between species (**Figure 4**), suggesting that they do subserve similar functions. Furthermore, a previous study by Kim and colleagues has measured the density of PV and VIP interneurons across mouse cortex using transgenic mouse lines (74). Using their data, we find a very similar gradient across mouse cortex when taking the ratio of PV and VIP interneurons (instead of PV and CR interneurons) (**Figure S5**).

Although a gradient in VIP interneurons was not considered in previous studies, Kim and colleagues did report a gradient in the density of PV relative to SST interneurons (74). This also seemed to indicate a general decrease of PV interneuron density towards higher cortical areas although a correlation with cortical hierarchy is less obvious, matching our findings with CB interneurons. The result of Kim and colleagues has been interpreted in terms of a gradient in the time constants of neural processing across the cortical hierarchy with activity becoming less transient and more sustained from sensory towards associatory areas (75). This interpretation also applies well to our findings, although we here add that a gradient in interneuron distribution could also provide a measure of the strength of feedforward versus feedback processing within the cortical hierarchy.

Our results challenge the currently prevalent view in human neuroscience that early visual areas are strongly influenced by top-down input, with a special role for V1 being at the bottom of the hierarchy, receiving a cascade of feedback input (9–11, 14, 15). Instead, our results suggest that early visual areas are dominated by bottom-up processing, in particular in V1.

One argument for the view that the processing in V1 is dominated by top-down input comes from retrograde tracing studies which indicated that 90% of the input comes from higher cortical areas, and with many more feedback than feedforward pathways targeting early visual areas (12). However, this is in terms of the number of neurons projecting to a target area, which is biased by the size of the source area and the proximity between the source and the target area. The feedforward input to V1 is a unique case as LGN is small and far from V1, making it hard to compare it’s contribution to nearby cortical areas. Our results allow a more direct comparison by using a marker of the type of neural processing intrinsic within areas.

Our results are in fact in line with retrograde tracing studies when excluding the LGN, with V2 for example receiving ∼ 75% of it’s external input from V1, and only ∼25% from higher cortical areas (61). Our results are also in line with a recent study that provided a sensitive measure of the density of thalamic axonal boutons in macaque monkey V1 (76), which found that it a factor 2 to 3 stronger than previously shown (27, 77). This suggest that the anatomical data that is used to constrain current computational models of the primate visual cortex would need to be updated significantly (78, 79).

An additional reason why early sensory areas have been considered to be strongly driven by top-down input is that studies using functional magnetic resonance imaging (fMRI) in humans have shown strong top-down modulation in early sensory areas (80), with examples of attentional modulation in V1 of more than 300% (81). However, fMRI only provides an indirect measure of neural activity, and has been suggested to be dominated by synaptic input in the superficial layers of the cortex where feedback input arrives (82). Direct neuronal recordings in primates suggest that top-down modulation in V1 is weak when compared to the feedforward driven response, generally around 5% (83), and maximally around 10% (84). And this modulation is known to gradually increase across the visual hierarchy, matching well with the gradient that we find in CR interneurons (56, 83, 85). Similarly strong attentional effects as measured with fMRI in humans have been found in macaque monkey V1 when using voltage sensitive imaging as measured on the surface of the cortex (86), indicating that a difference in top-down modulation in V1 is due to differences in recording methods, not to difference in animal species.

Our results are also in line with studies that have analyzed the distribution of cortical oscillatory activity in the brain, showing a gradient from lower to higher cortical areas with a decrease of the strength of gamma oscillations and a concurrent increase in strength of alpha / beta oscillations (87). Gamma oscillations have been implicated in fast feedforward processing (88, 89), and are linked to PV interneurons with fast time constants (33, 90, 91). In contrast, alpha and beta oscillations are implicated in more sluggish feedback processing (88, 89), and are linked to cell-type-specific microcircuits with longer time constants involving VIP and SST interneurons (90–92). Furthermore, a direct link has recently been found between the laminar distribution of cortical oscillations and interneuron distribution in macaque monkeys cortex (93).

We also found a clear gradient from the medial to lateral direction within macaque monkey vlPFC, matching a gradient that was previously found in anatomy and physiology. First of all, it matches classical anatomical studies on the connectivity pattern between the inferotemporal cortex (IT) and vlPFC, with middle IT projecting to area 8 and 45, and anterior IT projecting more laterally towards area 47/12 and orbitofrontal cortex (94, 95). Also, our results match with the three face patches that have been found along the medial to lateral axis in vlPFC, potentially matching the 3 pairs of face patches found in IT cortex (96, 97). Finally, our results match a recent study where electrical microstimulation in PFC was combined with whole-brain fMRI (98), finding that the posterior to anterior gradient in IT corresponded to a medial to lateral gradient in vlPFC.

Strong gradients can also be found in for example neurotransmitter receptors within the macaque monkey prefrontal cortex (99, 100), suggesting that it would be highly relevant to provide a more complete map within macaque monkey prefrontal cortex. One previous paper has already investigated interneuron distribution in a large number of areas within the macaque monkey prefrontal cortex (72). However, this paper counted neurons manually, limiting the data to a few tens of neurons per type and area. Also, different brain slices were used for different interneuron stainings, complicating a comparison between cell types even further, as large inhomogeneities in neuron density exist within an area. A recent study did collect a large dataset of PV and CR interneuron densities from dorsolateral PFC, as well as dorsal visual cortex (69). Their results show a similar increase of CR and decrease of PV from visual to prefrontal cortex, indicating that our results generalize to the dorsal pathway.

We found that the depth of the mouse cortical hierarchy is severely limited, corresponding in depth to early visual areas in primates (roughly V1 to V4). This result matches recent finding on the density of dendritic spines, showing a very shallow gradient in mice, corresponding to early sensory areas in primates (24). In contrast, the dendritic spine density in primate cortex continued to steeply increase towards higher areas, with only subtle differences between humans and macaque monkeys. Our results on a selective increase of CR interneurons in primates is also in line with a recent connectomics study, showing that disinhibitory circuits are selectively enhanced in both macaque monkeys and humans compared to mice (68). The strong overlap between humans and macaque monkeys also matches with the very limited genetic differences in the expression and distribution of cortical neurons in humans and non-human primates (45).

The method to use PV and CR or VIP as markers of feedforward versus feedback processing as proposed in the current study can be easily extended to other animal species, including humans, allowing a straight forward approach to quantitatively compare the depth and gradient of the cortical hierarchy across species.

Taken together, our results indicate that the mouse cortex remains driven by sensory information while the primate prefrontal cortex allows processing to become predominantly internal.

## Materials and Methods

### Animals

Two adult male macaque monkeys *(Macaca mulatta*), that had previously been used in other (non-invasive) protocols were used in this experiment. Three-month-old male mice, 20-30g, C57BL/6 (Janvier Laboratory) were used for the experiment (6 in total). Following their arrival in the laboratory, mice were kept under controlled conditions (ambient temperature of 20 to 25 °C) with a light/dark cycle of 12 hours on, 12 hours off, and with free access to food and water.

### Fixation and sectioning

All animals were euthanized and transcardially perfused with phosphate-buffered saline (PBS, 0.1M,, pH 7.4) followed by 4% paraformaldehyde (PFA) in PBS. After extraction, the brains were post-fixed in 4% PFA and then kept in PBS, at 4°C, until use. The macaque brains were manually sectioned into blocks corresponding to different brain areas of interest. All brains were transferred successively in 15% and 30% phosphate-buffered sucrose, at 4°C, until equilibrium, to prevent subsequent cryo-damage. Macaque brain blocks and mouse entire brains were shock-frozen and cut coronally into 20 µm-thick sections on a sliding freezing cryostat, where the sections were placed on cover glasses and stored at −80°C.

### Brain areas

For the delineation and terminology of brain areas we followed Petrides and Pandya 2007 for macaque monkeys (101), and the Allen Mouse Brain Common Coordinate Framework for mice (102), see also **Figure 2B** for a schematic. In macaque monkeys, we selected parts of V1 and V2 near the lunate sulcus and about 1cm lateral from the midline, as well as parts of V4 and TEO on opposite sides of the inferior occipital sulcus and close to the superior temporal sulcus (so all of them corresponding to parafoveal receptive fields (103, 104)). In mice, we selected primary visual cortex (Vp), lateromedial visual cortex (Vlm), anterolateral visual cortex (Val), posteromedial visual cortex (Vpm) and anteromedial visual cortex (Vam) in mouse cortex. In addition, we analyzed a set of areas in the ventrolateral prefrontal cortex (vlPFC) of monkeys which have recently been shown to lie on a functional gradient that matches the hierarchy in the ventral stream of the visual cortex (98), namely areas 8, 45 and 47/12 (parts c and o). These three areas also roughly correspond to the three frontal face patches : PA, PL and PO (96, 97). As a potential homologue, we analyzed three areas in the mouse medial frontal cortex (mFC) : prelimbic area (PL), infralimbic area (ILA) and medial orbital area (ORBm) (105). Finally, for both species we analyzed the anterior cingulate cortex (ACC, area 24a in macaque monkeys), which has been suggested to be relevant for cross-species comparison (106). We placed it at the top of the hierarchy based on electrophysiological results (107).

### Immunohistochemistry

All neurons and the three calcium-binding proteins (CR, CB and PV) were fluorescently tagged using specific antibodies (**Table S3**). The procedure was performed on multiple sections per area and per animal (**Table S2)**. Tissue sections were rehydrated in PBS, and then incubated in a blocking buffer comprising 1% IgG-free bovine serum albumin (BSA; Sigma A7030), 0.02% sodium azide (Sigma S2002), 1% Triton X-100 (Sigma X100), and 5% normal donkey serum (Eurobio CAEANE00-0U) in PBS for 2 hours at room temperature. A citrate step was performed by incubating sections in buffer comprising 10mM citrate sodium (Sigma W302600), and 0.5% Tween (Sigma P2287), 20 min at 85°C and then 15 min at room temperature. Sections were then transferred into fresh buffer comprising 1% IgG-free bovine serum albumin (BSA; Sigma A7030), 0.02% sodium azide (Sigma S2002), and 5% normal donkey serum (Eurobio CAEANE00-0U) in PBS with primary antibodies added. Sections were exposed to antibodies in a separate incubation steps. Antibodies against NeuN, CR and PV were combined on the same sections (in tissue from the 2 monkeys and 3 mice) and in parallel, antibodies against NeuN and CB were combined on other sections (in tissue from the same 2 monkeys and 3 additional mice). Tissue remained in the antibody solution for 1-2 hours at room temperature. After two rinsings with PBS, sections were incubated with their corresponding fluorescent secondary antibodies diluted in 1% IgG-free bovine serum albumin (BSA; Sigma A7030), 0.02% sodium azide (Sigma S2002), and 5% normal donkey serum (Eurobio CAEANE00-0U) in PBS for 1 hour at room temperature in the dark (separate incubation steps). All secondary antibodies were raised in donkey. Neuron-immunoreactive sites were always visualized with the Alexa 594 fluorophore (Alexa 594 donkey anti-mouse IgG, Abcam #150112, 1:250), and calcium-binding immunoreactive sites, CR, CB and PV were respectively visualized with the Alexa 488 fluorophore (Alexa 488 donkey anti-goat IgG, Abcam #150133, 1:250), the Alexa 350 fluorophore (Alexa 350 donkey anti-rabbit IgG, ThermoFisher #A10039, 1:250) and the Alexa 647 fluorophore (Alexa 647 donkey anti-guinea pig IgG, Jackson ImmunoResearch #706-605-148, 1:250) (**Table S3**). The sections were then rinsed in PBS, coverslipped with the ProLong Gold (PermaFluor TA-030-FM) cover medium and preserved in the dark at 4°C before imaging. To control for unspecific staining, we repeated the entire procedure but without the primary antibodies and found no fluorescent staining.

### Imaging and Image Processing

We used three or two filters to acquire fluorescent images of every section using Zeiss Axio Observer, at x10 magnification. We imaged entire sections for each animals. Images were captured using the Zeiss Zen 2 Blue edition software. For macaques, images were cropped and for each section the immunoreactive neurons were counted in two columns going perpendicularly from the pial surface to the white matter along the whole cortical depth. For mice, images were cropped to delineate the complete area. See **Figure S6** for examples of the selection of regions of interest in monkeys and mice. We used a previously established method to count cells (60). As a first step, we determined a luminance threshold to separate the signal pixels from the background pixels in each channel of each image. Then, two steps are applied in order to sequentially threshold each image, and to separate the aggregated particles (“Watershed” function of the ImageJ software). Cell bodies are finally selected using the “Analyze particles” function of the ImageJ software, applying a limit of size (from 20 to 500 pixels) and circularity (from 0.2 to 1) based on visual inspection of the results.

### Statistics

We used one-sample test of the regression line to test whether the slope of an interneuron gradient was different from zero (**Figure 3A, C**). For all tests involving two populations (**Figure S1**), we first used the F-test to test for equal variance of the populations. We subsequently used unpaired two-tailed t-tests and applied the Satterthwaite approximation in case variances were not equal.

## Acknowledgements

We thank Andreas Burkhalter for very helpful comments on an earlier version of the manuscript. The work was supported by a Young Investigator award from the French National Research Agency (grant agreement numbers : ANR-19-CE37-003-01) and a starting grant from the European Research Council under the European Union’s Horizon 2020 research and innovation program (grant agreement numbers : 101078667).

## Author contribution

Experimental design: M.G., F.G. and T.v.K. Data acquisition: M.G. with help from F.G. Data analysis: M.G. with help from T.v.K., Writing: M.G. and T.v.K. Supervision: TvK.

---

**Figure S1.**
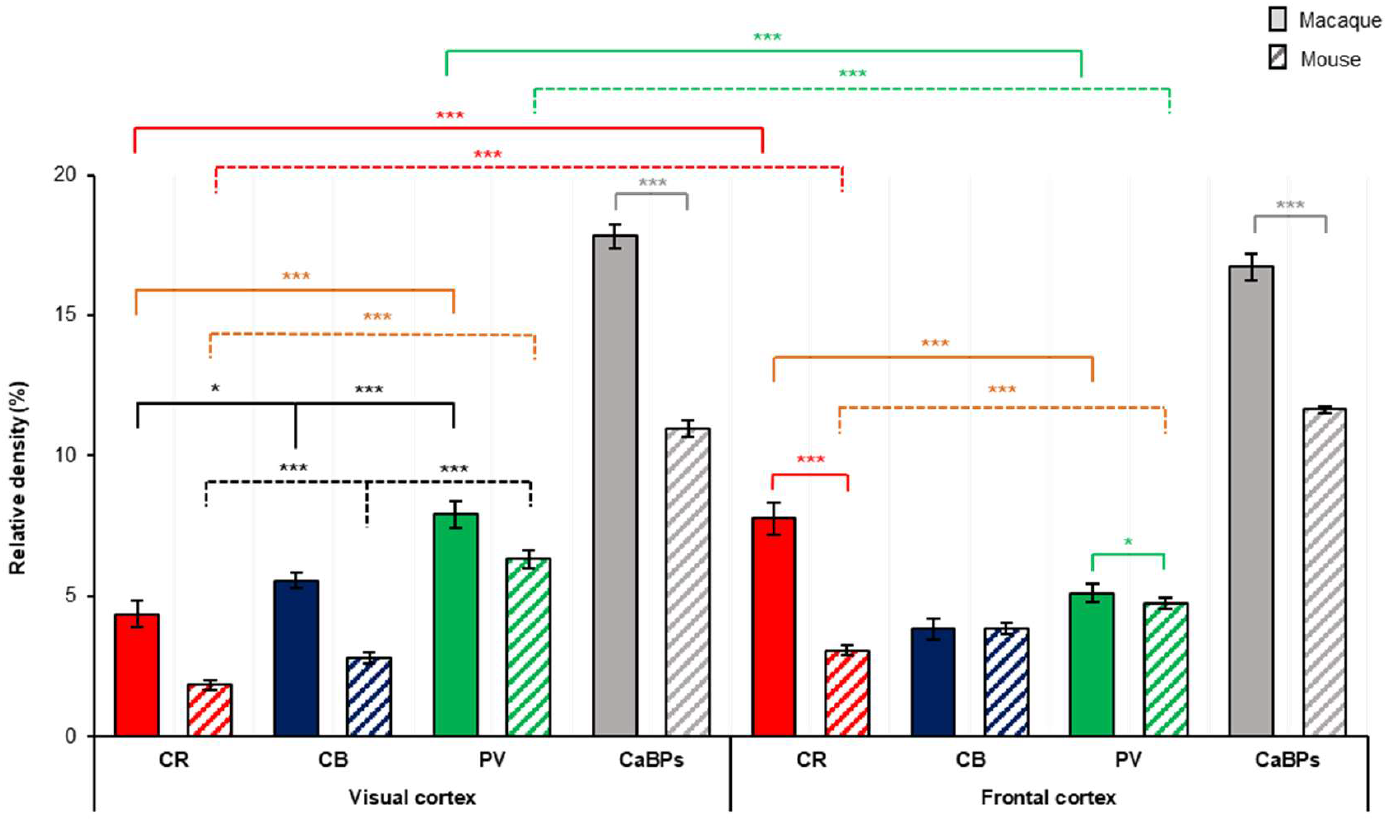
Calcium-binding interneuron densities for visual and frontal cortex in the macaque and mouse cortex. Distribution of the three interneurons cell types, CR (left column, red), CB (middle column, blue) and PV (right column, green), relative to the total neuronal population for macaque monkeys (homogenously colored) and mice (patterned). Error bars indicate standard error of the mean. \**P* < 0.01, \*\*\**P* < 0.0001

**Figure S2.**
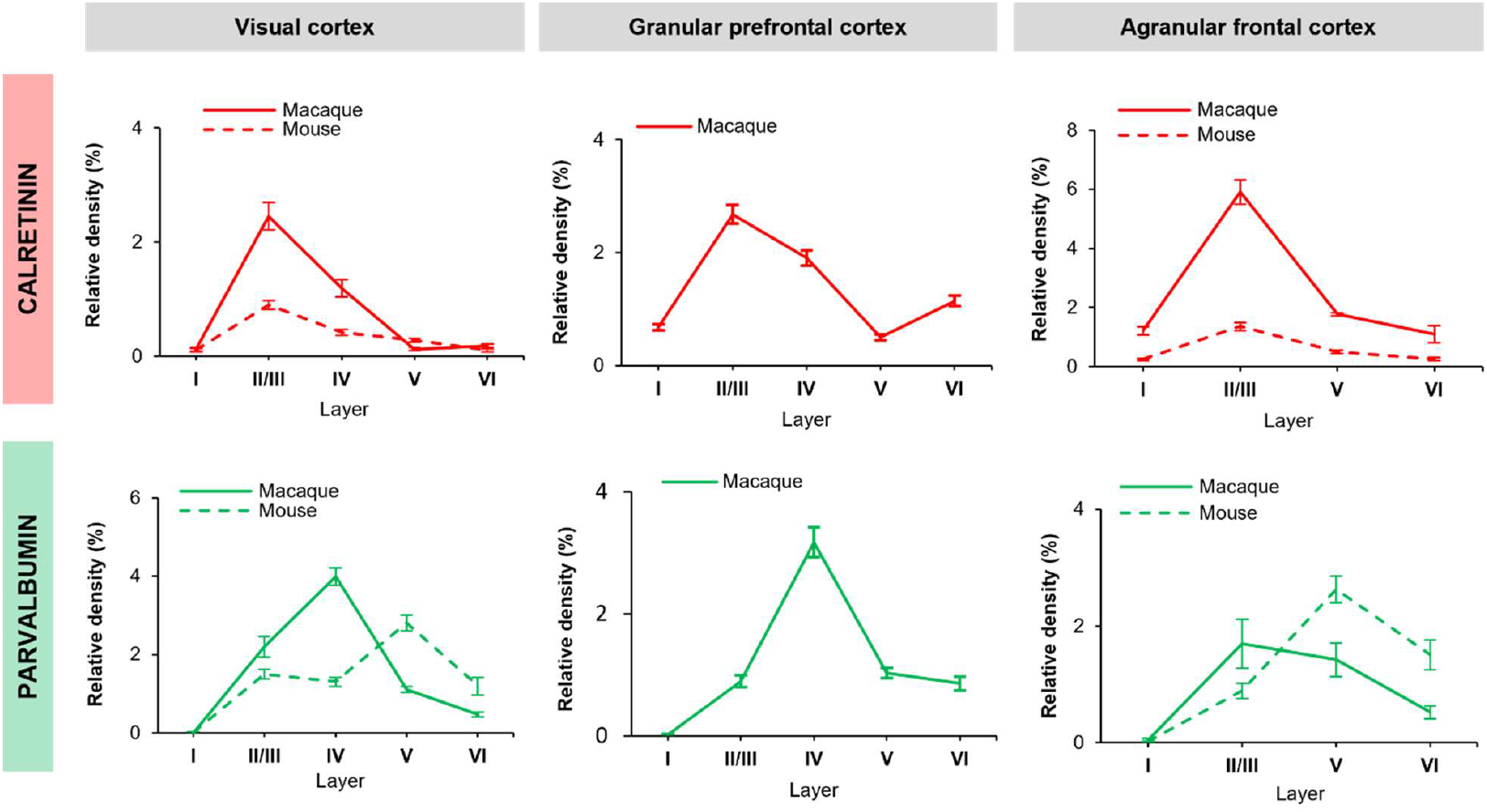
Laminar distribution of parvalbumin and calretinin interneurons. Densities of calretinin (top row) and parvalbumin (bottom row) interneurons relative to the total neuronal population across the depth of the visual (left column), granular prefrontal (middle column), and agranular prefrontal cortex (right column) in monkeys (continuous line) and mice (dashed line). Error bars indicate standard error of the mean.

**Figure S3.**
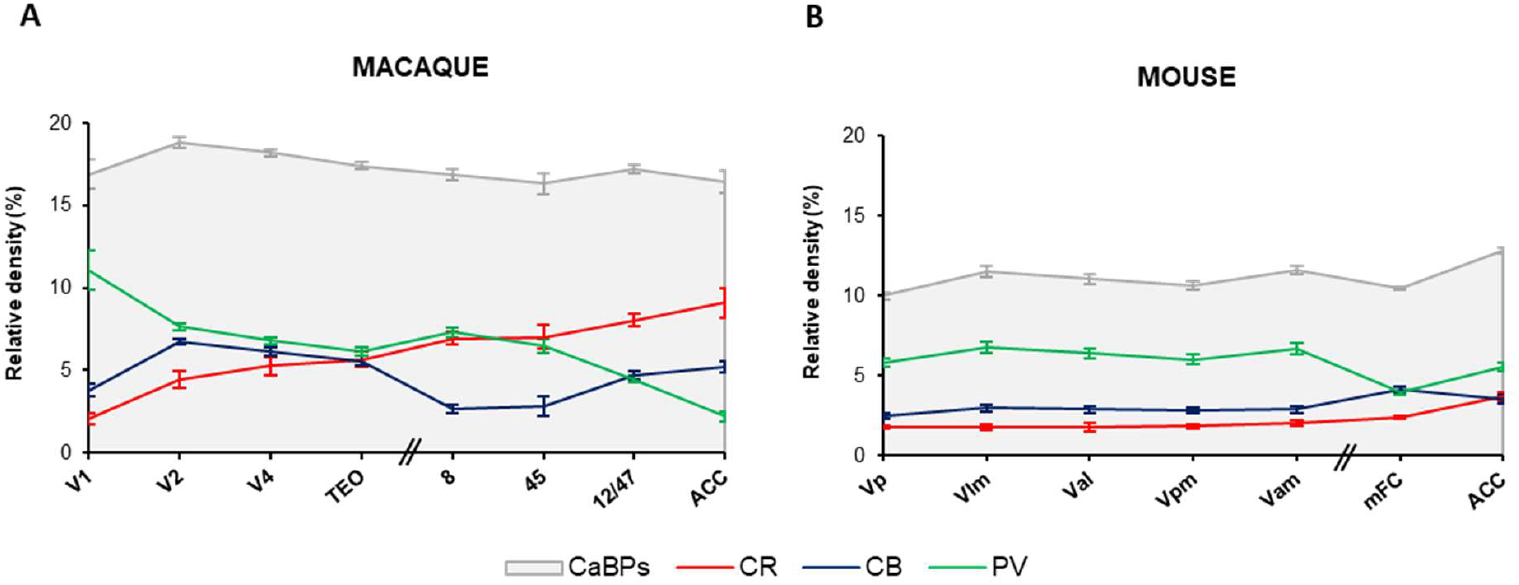
Calcium-binding interneuron distribution in the macaque and mouse cortex. Distribution of calretinin (CR, red), parvalbumin (PV, green) and calbindin (CB, blue) interneurons across the different areas in A) macaque and B) mouse cortex. With in grey, the sum of all three interneurons. Error bars indicate standard error of the mean.

**Figure S4.**
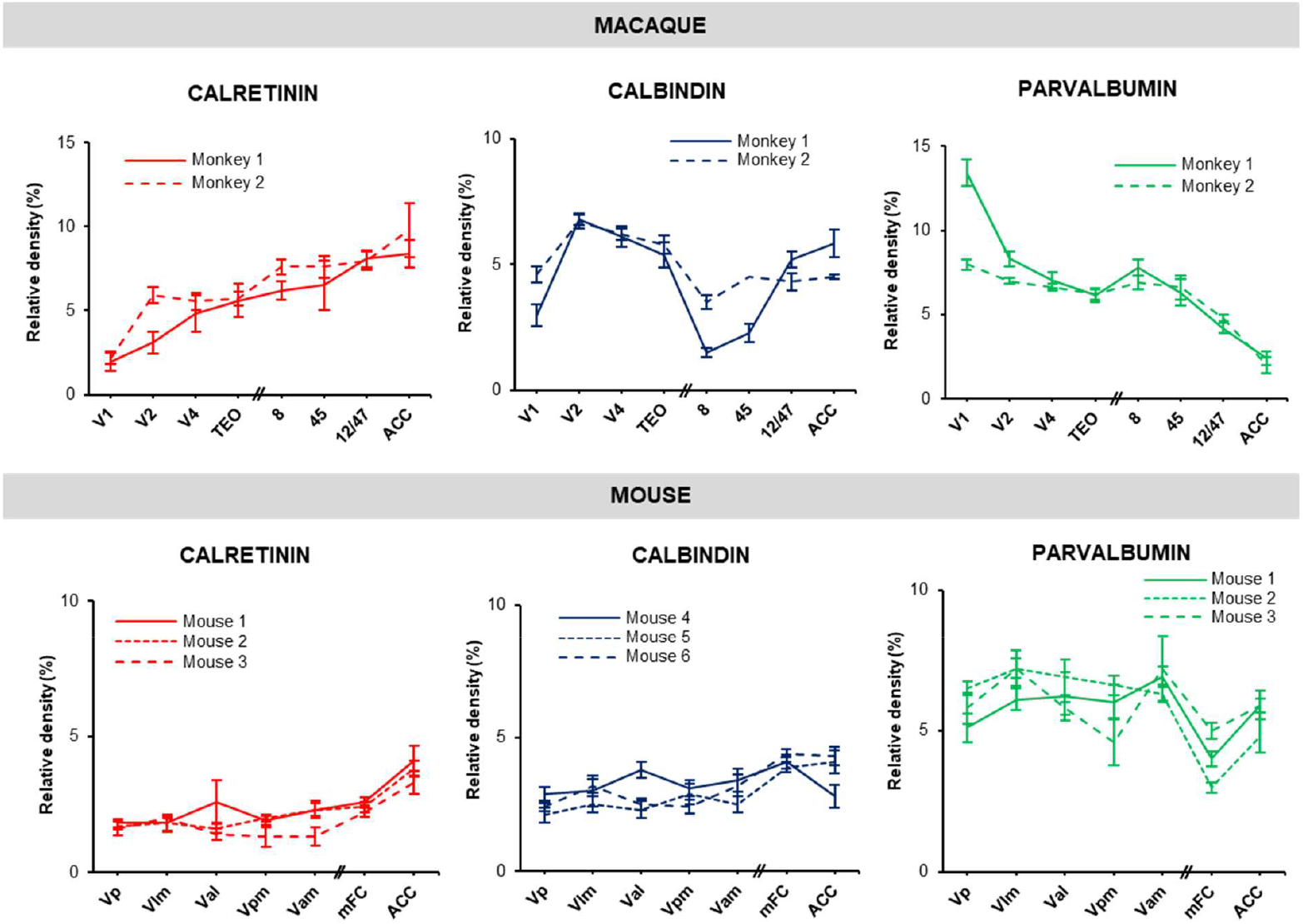
Calcium-binding interneuron distribution in the macaque and mouse cortex per individual. Distribution of three interneurons cell types, CR (left column, red), CB (middle column, blue) and PV (right column), relative to the total neuronal population for macaque monkeys (top row) and mice (bottom row), separately for each individual (continuous and dashed lines). Error bars indicate standard error of the mean.

**Figure S5.**
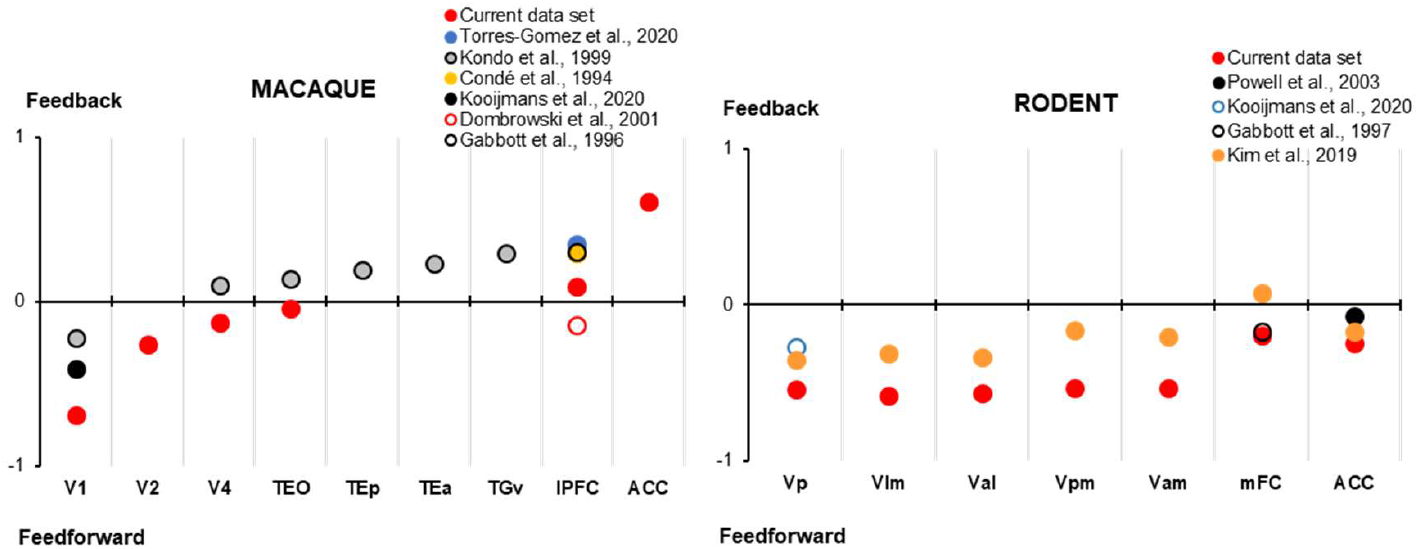
Normalized ratio of the interneurons involved in feedforward and feedback processing in the cortex of macaque monkeys and rodents, with a comparison to previous literature. The normalized difference between parvalbumin and calretinin interneurons is shown per area, calculated as (PV-CR)/(PV+CR). For Kim et al. 2019, VIP instead of CR interneurons are taken. For rodents, all results are from mice, except for Gabbott et al. 1997, where the results are from rats. The notation of the areas corresponds to the notation in the respective papers. Except for lateral PFC (lPFC) which corresponds to the average across all areas in ventro-and dorso-lateral PFC that are reported in the respective studies (i.e. areas 8-12 and / or 46), and ACC which corresponds to area 24a.

**Figure S6.**
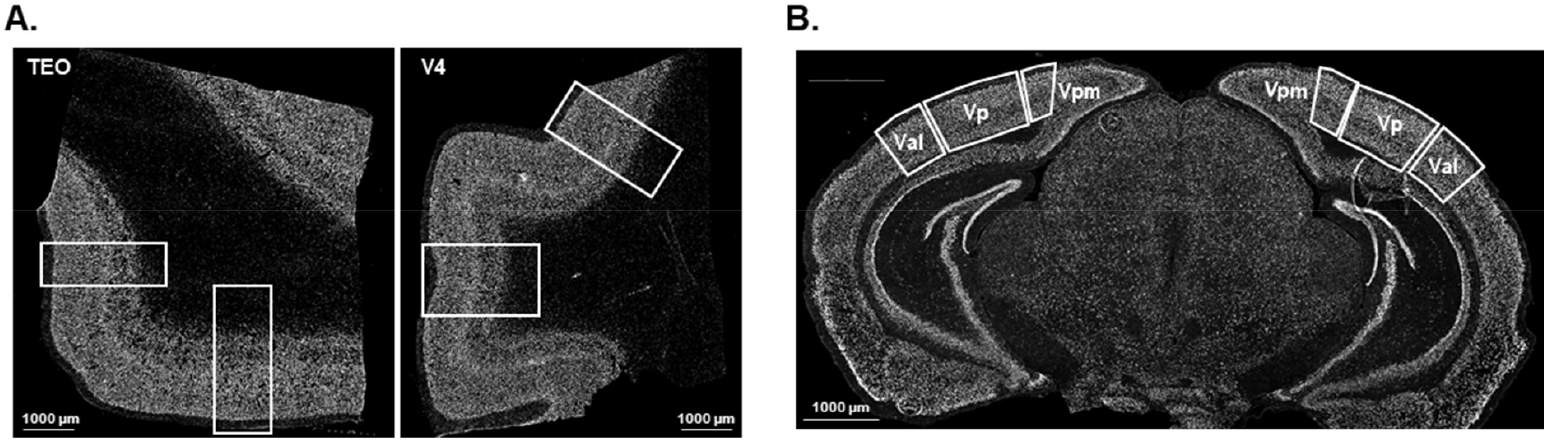
ROI delimitation in macaque monkey and mouse cortex. Representative examples of region of interest delimitation on specific A) monkey brain block slice, here TEO and V4 with two ROI, and on each entire B) mouse brain slice with several present areas (Vpm, Vp, Val).

**Table S1.** All tested antibodies in mouse and monkey cortex. With the name of the antibody, the name of the provider, the reference number of the antibody, the host species, and the results of the tests per species. A check mark indicates that the labeling worked, an x that it did not work, and a hyphen that this antibody was not tested in macaque monkeys (as it did not work in mice). The different dilutions that were used for the tests are indicated as well.

| Name | Provider | Ref | Host | Tested on |  |  |  |
| --- | --- | --- | --- | --- | --- | --- | --- |
|  |  |  |  | Mouse | Dilution | Macaque | Dilution |
| Anti-Calbindin | Abcam | ab82812 | Mouse | ✓ | 1/100 - 1/250 | ✓ | 1/100 |
| Anti-Calbindin | Swant | CB300 | Mouse | ✓ | 1/100 | ✓ | 1/1000 |
| Anti-Calbindin | Cliniscience | EB11759 | Goat | ✓ | 1/100 | ✓ | 1/500 |
| Anti-Calbindin D-28K | Swant | CB38 | Rabbit | ✓ | 1/250 | ✓ | 1/200 - 1/500 - 1/1500 |
| Anti-Calbindin D-28K | Thermo | PA5-50366 | Rabbit | ✓ | 1/100 | ✓ | 1/100 - 1/250 - 1/500 |
| Anti-Calbindin D9K | Santa Cruz | sc-74462 | Mouse | x | 1/100 | x | 1/50 - 1/250 - 1/500 - 1/1000 |
| Anti-Calbindin-D-28K | Sigma | C9848 | Mouse | ✓ | 1/100 | ✓ | 1/5000 |
| Anti-Calretinin | Swant | CG1 | Goat | ✓ | 1/50 - 1/100 | ✓ | 1/500 |
| Anti-Calretinin | Swant | CR7697 | Rabbit | ✓ | 1/100 | ✓ | 1/100 - 1/250 - 1/500 - 1/1000 - 1/2000 |
| Anti-Calretinin | Swant | CRgp7 | Guinea Pig | - |  | x | 1/100 - 1/500 - 1/1000 - 1/2000 |
| Anti-Calretinin / CALB2 | Cliniscience / LSBio | C-150-348 | Sheep | ✓ | 1/100 | x | 1/250 - 1/500 - 1/1000 |
| Anti-CamKII delta | Thermo | PA5-14036 | Rabbit | - |  | x | 1/50 - 1/250 - 1/500 - 1/1000 - 1/2000 |
| Anti-CamKII gamma | Abcam | ab37999 | Rabbit | - |  | x | 1/25 - 1/50 - 1/100 |
| Anti-GABA | Sigma | A0310 | Mouse | ✓ | 1/50 - 1/100 - 1/500 | x | 1/50 - 1/100 - 1/500 |
| Anti-GAD65+GAD67 | Abcam | ab11070 | Rabbit | - |  | x | 1/200 - 1/1000 - 1/2000 |
| Anti-GFAP | Abcam | ab53554 | Goat | ✓ |  | ✓ | 1/500 - 1/750 |
| Anti-HSP70/HSP72 | Enzo | ADI-SPA-810 | Mouse | - |  | x | 1/25 - 1/50 - 1/100 |
| Anti-Iba1 | Abcam | ab107159 | Goat | ✓ | 1/500 | x | 1/250 - 1/500 - 1/1000 - 1/1500 - 1/2000 |
| Anti-Iba1 | Wako | 019-19741 | Rabbit | ✓ | 1/100 | ✓ | 1/100 - 1/250 - 1/500 - 1/1000 - 1/1500 |
| Anti-NeuN | Abcam | ab177487 | Rabbit | ✓ | 1/1000 | ✓ | 1/1000 |
| Anti-NeuN | Millipore / Chemicon | MAB377 | Mouse | ✓ | 1/100 | ✓ | 1/100 - 1/500 - 1/1000 |
| Anti-Parvalbumin | Abcam | ab32895 | Goat | ✓ | 1/100 | ✓ | 1/100 |
| Anti-Parvalbumin | Swant | PVG213 | Goat | ✓ | 1/500 - 1/1500 - 1/3000 | ✓ | 1/500 - 1/1500 - 1/3000 |
| Anti-Parvalbumin | Swant | PV27 | Rabbit | ✓ |  | ✓ | 1/500 - 1/1000 - 1/2500 |
| Anti-Parvalbumin | Swant | GP72 | Guinea Pig | ✓ | 1/100 | ✓ | 1/500 - 1/1000 - 1/2500 |
| Anti-Parvalbumin | Abcam | ab181086 | Rabbit | ✓ | 1/100 | ✓ | 1/100 - 1/250 - 1/500 - 1/750 - 1/1000 - 1/2000 |
| Anti-Parvalbumin | Thermo | PA5-47693 | Sheep | ✓ | 1/1000 | ✓ | 1/100 - 1/250 - 1/500 - 1/1000 - 1/1500 |
| Anti-Parvalbumin | Swant | PV 235 | Mouse | ✓ | 1/5000 - 1/10000 - 1/20000 | ✓ | 1/5000 - 1/10000 - 1/20000 |
| Anti-Somatostatin | Enzo | BML-SA1267 | Rabbit | ✓ | 1/100 | x | 1/50 - 1/100 - 1/500 |
| Anti-Somatostatin | NeoBiotech / Cliniscience | NB-22-14395 | Rabbit | x | 1/100 | x | 1/100 - 1/250 - 1/500 |
| Anti-Somatostatin | Peninsula / BMA | T-4103 | Rabbit | ✓ | 1/100 | x | 1/50 - 1/100 - 1/250 - 1/500 |
| Anti-Somatostatin | Bioss | bs-8877R | Rabbit | - |  | x | 1/100 - 1/500 |
| Anti-Somatostatin | Thermo | MA5-16987 | Rat | - |  | x | 1/50 - 1/100 |
| Anti-Somatostatin | Proteogenix | PTX12661 | Rabbit | ✓ | 1/100 | x | 1/100 - 1/500 - 1/1000 |
| Anti-Somatostatin | Millipore | MAB354 | Rat | - |  | x | 1/100 |
| Anti-Somatostatin | Atlas Antibodies | HPA019472 | Rabbit | ✓ | 1/100 | x | 1/100 |
| Anti-Somatostatin | ABclonal | A9274 | Rabbit | - |  | x | 1/100 |
| Anti-Somatostatin | MyBioSource | MBS536200 | Sheep | - |  | x | 1/100 |
| Anti-Somatostatin | MyBioSource | MBS822270 | Rabbit | - |  | x | 1/100 |
| Anti-Somatostatin | Abcam | ab30788 | Rat | ✓ | 1/100 | x | 1/25 - 1/50 - 1/100 - 1/250 - 1/500 |
| Anti-Somatostatin | Abcam | ab64053 | Rabbit | ✓ | 1/100 | x | 1/100 - 1/250 |
| Anti-VIP | Immunostar | 20077 | Rabbit | x | 1/500 - 1/1500 - 1/3000 - 1/5000 - 1/8000 - 1/10000 | x | 1/500 - 1/1500 - 1/3000 - 1/5000 - 1/8000 - 1/10000 |

**Table S2.** Total number of sections analysed. Number of counted A) slices of mouse brain, and B) regions of macaque brain, of each areas in each animals.

|  | CR/PV |  |  |  | CB |  |  |  |
| --- | --- | --- | --- | --- | --- | --- | --- | --- |
|  | Mouse 1 | Mouse 2 | Mouse 3 | Total | Mouse 4 | Mouse 5 | Mouse 6 | Total |
| VISp | 12 | 12 | 6 | 30 | 6 | 6 | 9 | 21 |
| VISIm | 6 | 8 | 2 | 16 | 6 | 5 | 7 | 18 |
| VISal | 3 | 5 | 4 | 12 | 6 | 6 | 6 | 18 |
| VISpm | 6 | 7 | 3 | 16 | 6 | 6 | 6 | 18 |
| VISam | 6 | 12 | 6 | 24 | 6 | 11 | 4 | 21 |
| mFC | 33 | 35 | 25 | 93 | 30 | 18 | 19 | 67 |
| ACC | 15 | 14 | 14 | 43 | 11 | 7 | 8 | 26 |

|  | CR/PV |  |  | CB |  |  |
| --- | --- | --- | --- | --- | --- | --- |
|  | Monkey 1 | Monkey 2 | Total | Monkey 1 | Monkey 2 | Total |
| V1 | 4 | 3 | 7 | 5 | 4 | 9 |
| V2 | 6 | 6 | 12 | 7 | 5 | 12 |
| V4 | 5 | 7 | 12 | 5 | 3 | 8 |
| TEO | 5 | 6 | 11 | 5 | 5 | 10 |
| 8 | 8 | 10 | 18 | 8 | 10 | 18 |
| 45 | 2 | 2 | 4 | 3 | 1 | 4 |
| 12/47 | 12 | 10 | 22 | 10 | 12 | 22 |
| ACC | 4 | 4 | 8 | 4 | 4 | 8 |

**Table S3.**
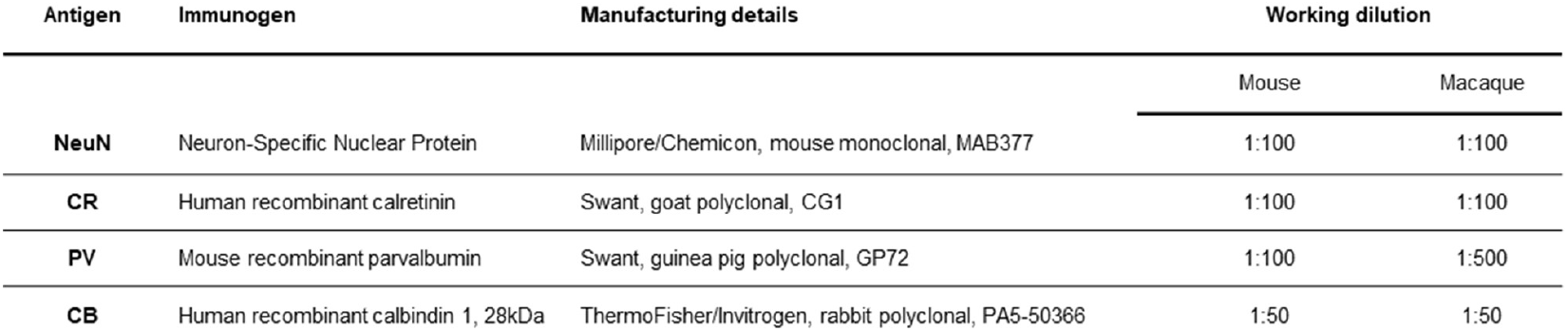
Detailed information for the primary antibodies used for immunostaining.

| Antigen | Immunogen | Manufacturing details | Working dilution |  |
| --- | --- | --- | --- | --- |
|  |  |  | Mouse | Macaque |
| NeuN | Neuron-Specific Nuclear Protein | Millipore/Chemicon, mouse monoclonal, MAB377 | 1:100 | 1:100 |
| CR | Human recombinant calretinin | Swant, goat polyclonal, CG1 | 1:100 | 1:100 |
| PV | Mouse recombinant parvalbumin | Swant, guinea pig polyclonal, GP72 | 1:100 | 1:500 |
| CB | Human recombinant calbindin 1, 28kDa | ThermoFisher/Invitrogen, rabbit polyclonal, PA5-50366 | 1:50 | 1:50 |

